# Unc 51–like autophagy-activating kinase (ULK1) mediates clearance of free α-globin in β-thalassemia

**DOI:** 10.1101/454652

**Authors:** Christophe Lechauve, Julia Keith, Eugene Khandros, Stephanie Fowler, Kalin Mayberry, Abdullah Freiwan, Christopher S. Thom, Paola Delbini, Emilio Boada Romero, Jingjing Zhang, Irene Motta, Heather Tillman, M. Domenica Cappellini, Mondira Kundu, Mitchell J. Weiss

## Abstract

Erythroid maturation is coordinated to maximize the production of hemoglobin A heterotetramers (α2β2) and minimize the accumulation of potentially toxic free α- or β-globin subunits. In β-thalassemia, mutations in the β-globin gene cause a build-up of free α-globin, which forms intracellular precipitates that impair erythroid cell maturation and viability. Protein quality-control systems mitigate β-thalassemia pathophysiology by degrading toxic free α-globin. We show that loss of the Unc 51–like autophagy-activating kinase gene *Ulk1* in β-thalassemic mice reduces autophagic clearance of α-globin in red cell precursors and exacerbates disease phenotypes, whereas inactivation of the canonical autophagy gene *Atg5* has minimal effects. Systemic treatment with rapamycin to inhibit the ULK1 inhibitor mTORC1 reduces α-globin precipitates and lessens pathologies in β-thalassemic mice, but not in those lacking *Ulk1*. Similarly, rapamycin reduces free α-globin accumulation in erythroblasts derived from β-thalassemic patient CD34^+^ hematopoietic progenitors. Our findings identify a new, drug-regulatable pathway for ameliorating β-thalassemia.

**One Sentence Summary:** Rapamycin alleviates β-thalassemia by stimulating ULK1-dependent autophagy of toxic free α-globin.

## Main Text

β-Thalassemia is a common, frequently debilitating inherited anemia caused by *HBB* gene mutations that reduce or eliminate expression of the β-globin subunit of adult hemoglobin (HbA, α2β2)(*1*). The pathophysiology arises from two major consequences of β-globin deficiency. First, insufficient HbA production reduces red blood cell (RBC) oxygen-carrying capacity, causing tissue hypoxia. Second, unstable free α-globin protein generates cytotoxic reactive oxidant species and cellular precipitates that impair the maturation and viability of RBC precursors (resulting in ineffective erythropoiesis) and induce premature lysis of mature RBCs (termed hemolysis)(*2–9*). Heterozygosity for β-globin null mutations (β-thalassemia trait) is usually clinically silent but becomes symptomatic with co-inheritance of extra α-globin genes (*HBA1* or *HBA2*), which increases the level of free α-globin protein(*10–13*). Hence, globin-chain imbalance with accumulation of free α-globin is a primary determinant of β-thalassemia pathophysiology.

Normal and β-thalassemic RBC precursors have mechanisms to detoxify excess α-globin (reviewed in reference ^(*11*)^). Specifically, free α-globin is stabilized by the molecular chaperone alpha hemoglobin stabilizing protein (AHSP)(*14, 15*) and is eliminated by other protein quality-control (PQC) mechanisms, including the ubiquitin-proteasome system and autophagy(*9*). Together, these observations suggest that individuals with β-thalassemia mutations can tolerate a modest pool of free α-globin, with pathologies arising only when the level of free α-globin exceeds the detoxification capacity of the endogenous protective mechanisms (reviewed in reference ^(*11*)^). In this respect, β-thalassemia resembles “protein aggregation disorders,” characterized by the accumulation of various toxic protein precipitates in disease-specific target tissues (reviewed in references (*9, 16–19*)). Efforts are underway to treat protein aggregation disorders via the pharmacologic induction of PQC pathways to eliminate relevant unstable proteins(*20–23*). In the current study, we sought to better define the mechanisms of free α-globin clearance by autophagy and to manipulate this pathway in order to treat β-thalassemia.

Autophagy is a process that delivers cytoplasmic proteins or organelles to lysosomes for degradation(*24*). Numerous forms of autophagy are essential for eukaryotic tissue development, cellular homeostasis, and protection against various stresses(*25–28*). The major autophagic pathway, termed “macroautophagy” (hereafter referred to as “autophagy”), is a multistep process that envelops cargo to be degraded within a double-membrane vesicle (an autophagosome) that eventually fuses with lysosomes (reviewed in references(*24, 29–31*)). Research by several groups, including ours, indicates that free α-globin is degraded by autophagy in β-thalassemia (*9, 20, 32*). Here we examined the contributions of two core autophagy proteins, Unc-51–like kinase 1 (ULK1) and ATG5, to disease progression in a mouse model of β-thalassemia. ULK1 initiates autophagosome formation by phosphorylating several autophagy proteins in response to cellular stresses and has additional roles in autophagy-independent vesicular transport(*33, 34*). ATG5 is part of a protein complex that conjugates microtubule-associated protein 1 light chain 3 beta (Map1LC3b, henceforth referred to as LC3) and related proteins to membrane phosphatidyl serine, creating a scaffold that facilitates cargo recognition and autophagosome maturation(*21, 24*). ULK1 and ATG5 are essential for most, but not all, autophagy processes. For example, during RBC formation, ULK1 has a non-redundant role in eliminating mitochondria and ribosomes from reticulocytes, whereas ATG5 is dispensable for these processes(*35–37*).

We used gene-targeted mice to show that the accumulation of toxic insoluble α-globin is inhibited by autophagy that is largely ULK1-mediated and ATG5-independent. Moreover, activation of ULK1 by the mTORC1 (mechanistic Target Of Rapamycin Complex 1) inhibitor rapamycin reduced the accumulation of insoluble free α-globin and alleviated β-thalassemic phenotypes in mice and in human patient–derived RBC precursors. Our findings provide definitive evidence that autophagy relieves the effects of β-thalassemia mutations by degrading excess α-globin, and identify an associated mechanism that can be targeted to develop new therapies.

### Hematopoietic ablation of *Ulk1* exacerbates β-thalassemia

To determine whether ULK1 mediates free α-globin degradation in β-thalassemia, we introduced a null allele into β-thalassemic mice (strain *Hbb*^*Th3*/+^)(*38, 39*). Double-mutant *Hbb*^*Th3*/+^*Ulk1*^−/−^ mice were born at a normal Mendelian ratio but exhibited increased perinatal death. Only 15% of *Hbb*^*Th3*/+^*Ulk1*^−/−^ mice survived for 30 days, compared to 80% of *Hbb*^*Th3*/+^*Ulk1*^+/+^ mice (*P* < 0.01) (Fig. 1A). To examine the hematopoietic cell−intrinsic consequences of ULK1 loss in β-thalassemia, we transplanted embryonic (E) day 14.5 fetal liver cells from *Hbb*^*Th3*/+^*Ulk1*^−/−^ and control strains (CD45.2) into lethally irradiated wild-type (WT) CD45.1 hosts (Fig. 1B). Donor hematopoietic cell engraftment exceeded 92% after 90 days (Fig. S1A). Loss of *Ulk1* exacerbated β-thalassemia, with a 20% reduction in the RBC count (*P* < 0.0001), and a 2-fold increase in the reticulocyte count (*P* < 0.0001) to maintain the same level of blood hemoglobin (Hb) (Fig. 1C–E and Table S1). Biotin labeling experiments showed that loss of *Ulk1* reduced the half-life of circulating RBCs from 7.5 days (for *Hbb*^*Th3*/+^*Ulk1*^+/+^ RBCs) to 5.4 days (for *Hbb*^*Th3*/+^*Ulk1*^−/−^ RBCs) (*P* < 0.0001), as compared to 17.5 days for WT (*Hbb^+/+^Ulk1^+/+^*) RBCs (Fig. 1F). Loss of ULK1 in *Hbb*^*Th3*/+^ mice also caused increased splenomegaly, bone marrow erythroid precursor hyperplasia, and extramedullary erythropoiesis in spleen and liver (Fig. 1G–H, Fig. S1B–E, and Table S2). Together, these findings show that ULK1 deficiency exacerbates ineffective erythropoiesis and reduces the viability of mature RBCs.

**Fig. 1.**
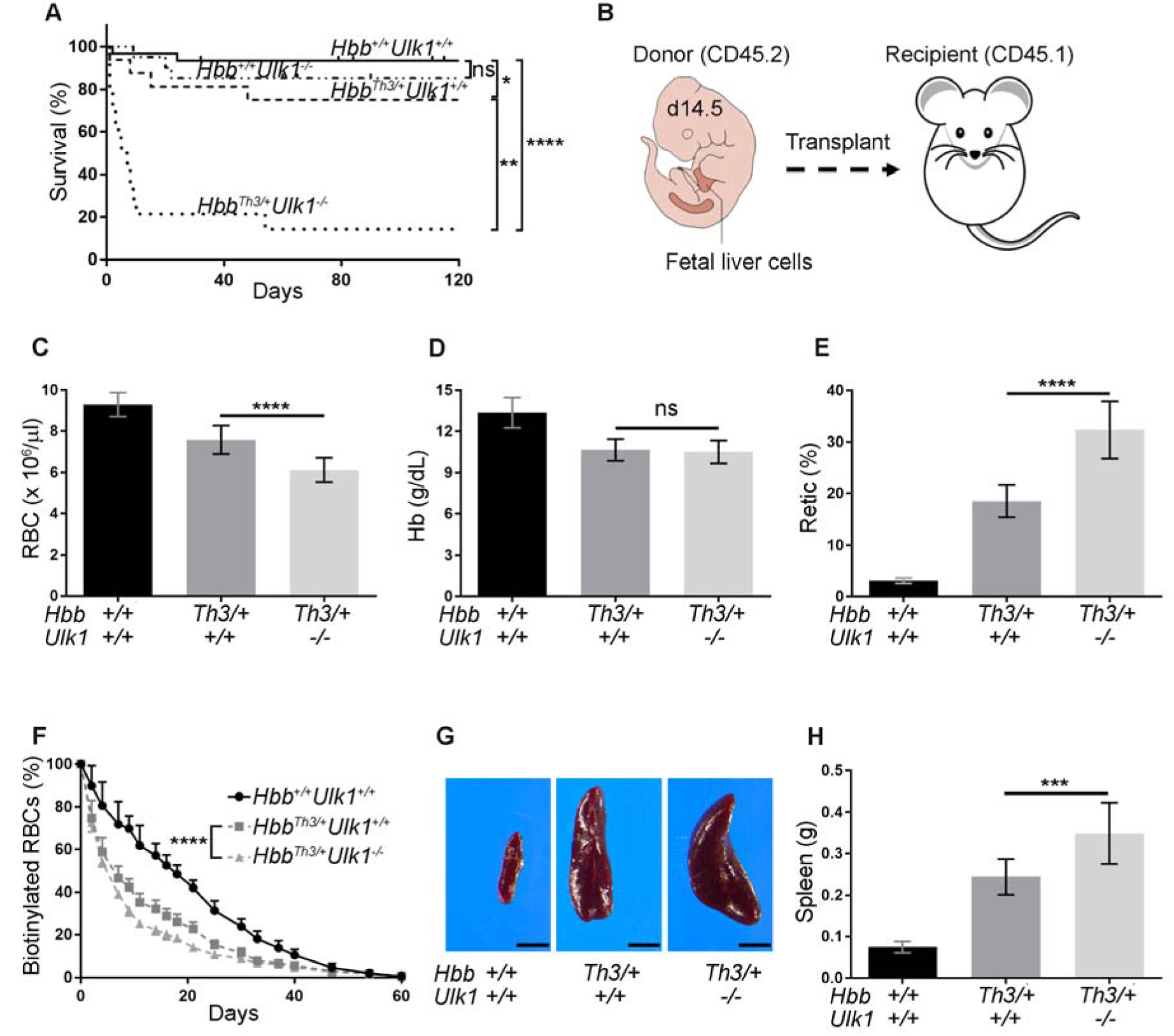
Hematopoietic cell ablation of *Ulk1* exacerbates β-thalassemia. (**A**) Kaplan-Meier survival plot of mice with genotypes *Hbb^+/+^Ulk1^+/+^*, n = 30; *Hbb^+/+^Ulk1^−/−^*, n = 20; *Ulk1*^+/+^*Hbb*^*Th3*/+^, n = 20; and *Hbb*^*Th3*/+^*Ulk1*^−/−^, n = 15. (**B**) Fetal liver hematopoietic stem cell (HSC) transplantation. Embryonic (E) day 14.5 fetal liver cells with the genotypes shown in panel a (C57BL/6, CD45.2) were injected intravenously into lethally irradiated WT mice (C57BL/6, CD45.1). (**C–E**) Erythroid indices (y-axis) according to donor HSC genotype (x-axis) at 90 days after HSC transplant. WT, n = 22; *Hbb*^*Th3*/+^*Ulk1*^+/+^, n = 29; *Hbb*^*Th3*/+^*Ulk1*^−/−^, n = 25. (**F**) Red blood cell (RBC) survival. At time 0, sulfo-NHS-biotin was injected into the tail vein of mice at 45 days post transplant with HSCs of the indicated genotypes. The fraction of biotinylated RBCs over time was quantified by streptavidin labeling followed by flow cytometry. The measured RBC half-lives for the genotypes are 17.5 days for *Hbb^+/+^Ulk1^+/+^*, n = 8; 7.5 days for *Hbb*^*Th3*/+^*Ulk1*^+/+^, n = 7; and 5.4 days for *Hbb*^*Th3*/+^*Ulk1*^−/−^, n = 4. (**G**) Representative spleens from mice with HSC donors of the indicated genotypes. (**H**) Summary of spleen weights: *Hbb^+/+^Ulk1^+/+^*, n = 19; *Hbb*^*Th3*/+^*Ulk1*^+/+^, n = 15; *Hbb*^*Th3*/+^*Ulk1*^−/−^, n = 12. All graphs show the mean ± SD; \*\*\*\**P* < 0.0001; \*\*\**P* < 0.001; \*\**P* < 0.01; \**P* < 0.05; ns: not significant.

### ULK1 loss reduces clearance of free α-globin in β-thalassemia

To investigate whether the loss of ULK1-dependent autophagy caused increased accumulation of precipitated α-globin, we lysed circulating RBCs and fractionated the contents by centrifugation followed by triton–acetic acid–urea (TAU) gel electrophoresis to resolve the α- and β-globin proteins(*9*). As expected, *Hbb*^*Th3*/+^ RBCs contained insoluble α-globin, which was increased approximately 2-fold by the loss of ULK1 (*P* < 0.001) (Fig. 2A,B). Transmission electron microscopy (TEM) revealed collections of electron-dense material, presumably α-globin precipitates, in flow cytometry–purified reticulocytes from β-thalassemic mice, but not in those from control mice (Fig. 2C). Automated image analysis of 100 to 200 cells from multiple mice demonstrated an approximately 2-fold increased area of electron-dense material in *Hbb*^*Th3*/+^*Ulk1*^−/−^ reticulocytes as compared to *Hbb*^*Th3*/+^*Ulk1*^+/+^ reticulocytes (*P* < 0.05) (Fig. 2D). Moreover, the levels of the autophagy adaptor protein P62 and lipidated forms of the autophagosomal marker protein LC3 were increased in reticulocytes of *Hbb*^*Th3*/+^ mice, as compared to WT (*Hbb^+/+^Ulk1^+/+^*) reticulocytes, and were even higher in Hbb^Th3/+^Ulk1^−/−^ mice (Fig. 2E). Thus, the loss of ULK1 increases the accumulation of insoluble α-globin and autophagic compartments in β-thalassemia.

**Fig. 2.**
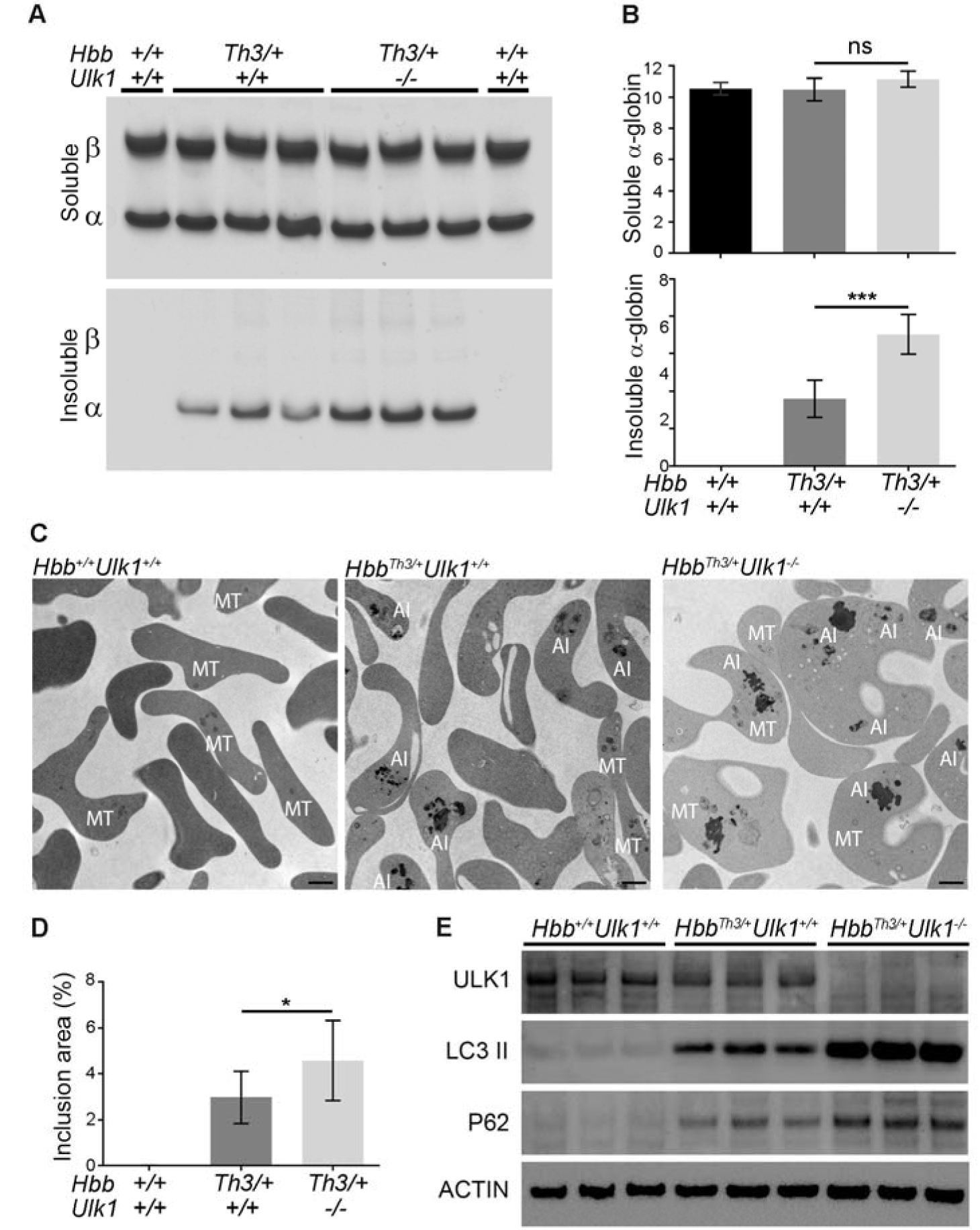
ULK1 loss in β-thalassemia causes increased accumulation of insoluble free α-globin. Wild-type mice were transplanted with fetal liver HSCs as in Figure 1 and analyzed 90–120 days later. (**A**) Soluble and insoluble globins in RBCs. Equal numbers of RBCs (from approximately 20 µL of blood, adjusted on hematocrit percentage) were lysed, centrifuged to separate the insoluble and soluble proteins, fractionated by triton– acetic acid–urea (TAU) gel electrophoresis to resolve the α- and β-globin chains, and stained with Coomassie brilliant blue. (**B**) Results of multiple experiments performed as in panel E. The y-axis shows the relative globin staining intensity on TAU gels, measured by automated image analysis. *Hbb^+/+^Ulk1^+/+^*, n = 4; *Hbb*^*Th3*/+^*Ulk1*^+/+^, n = 8; *Hbb*^*Th3*/+^*Ulk1*^−/−^, n = 8. (**C**) Purified reticulocytes analyzed by transmission electron microscopy (TEM). AI, α-globin inclusions; MT, mitochondria. Scale bars represent 1 µm. (**D**) Areas of electron-dense α-globin inclusions in reticulocytes (y-axis) were quantified by automated image analysis of electron micrographs. *Hbb^+/+^Ulk1^+/+^*, n = 7; *Hbb*^*Th3*/+^*Ulk1*^+/+^, n = 11; *Hbb*^*Th3*/+^*Ulk1*^−/−^, n = 12. A total of 100 to 200 cells from each mouse were analyzed. (**E**) Western blot analysis for autophagic markers of reticulocytes from mice of indicated genotypes. All graphs show the mean ± SD; \*\*\**P* < 0.001; \**P* < 0.05; ns: not significant.

### ATG5-independent autophagy of α-globin clearance in β-thalassemia

ATG5 is required for most autophagy processes. However, mitochondria are eliminated from reticulocytes in an ATG5-independent, ULK1-dependent form of autophagy(*35–37*). Therefore, we investigated the role of ATG5 in eliminating free α-globin. We introduced into *Hbb*^*Th3*/+^ β-thalassemic mice a conditional *Atg5* allele, in which exon 3 is flanked by loxP sites(*26*), and a Cre recombinase transgene (*EpoR*-*Cre*) that is expressed exclusively in erythroid precursors(*40*) (Fig. S2A). Double-mutant (*Hbb*^*Th3*/+^*Atg5*^*fl/fl*^) mice were born at a normal Mendelian ratio (data not shown). At age 4–6 months, *Atg5* transcripts and protein levels were reduced by 90% to 95% in Ter119^+^ bone marrow erythroid precursors (Fig. S2B). The erythroid indices of *Hbb*^*Th3*/+^*Atg5*^*fl/fl*^ mice were mildly worsened compared to those of *Hbb*^*Th3*/+^ mice, as reflected by 10% reductions in the RBC counts (*P* < 0.001) and 20% increases in the reticulocyte counts (*P* < 0.05) (Fig. 3A–C and Table S3). ATG5 deficiency in *Hbb*^*Th3*/+^ mice had minimal effect on spleen weight (Fig. 3D). Consistent with the expected impairment in autophagy (reviewed in (*41*)), the levels of the autophagy adaptor protein P62 were increased in RBC insoluble fractions of *Atg5*^*fl/fl*^ and *Hbb*^*Th3*/+^*Atg5*^*fl/fl*^ mice (Fig. S2C). However, ATG5 deficiency in *Hbb*^*Th3*/+^ mice had minimal effects on the levels of insoluble α-globin (Fig. 3E,F). Thus, ATG5 mediates some autophagy processes during erythropoiesis, but is not required for the elimination of free α-globin.

**Fig. 3.**
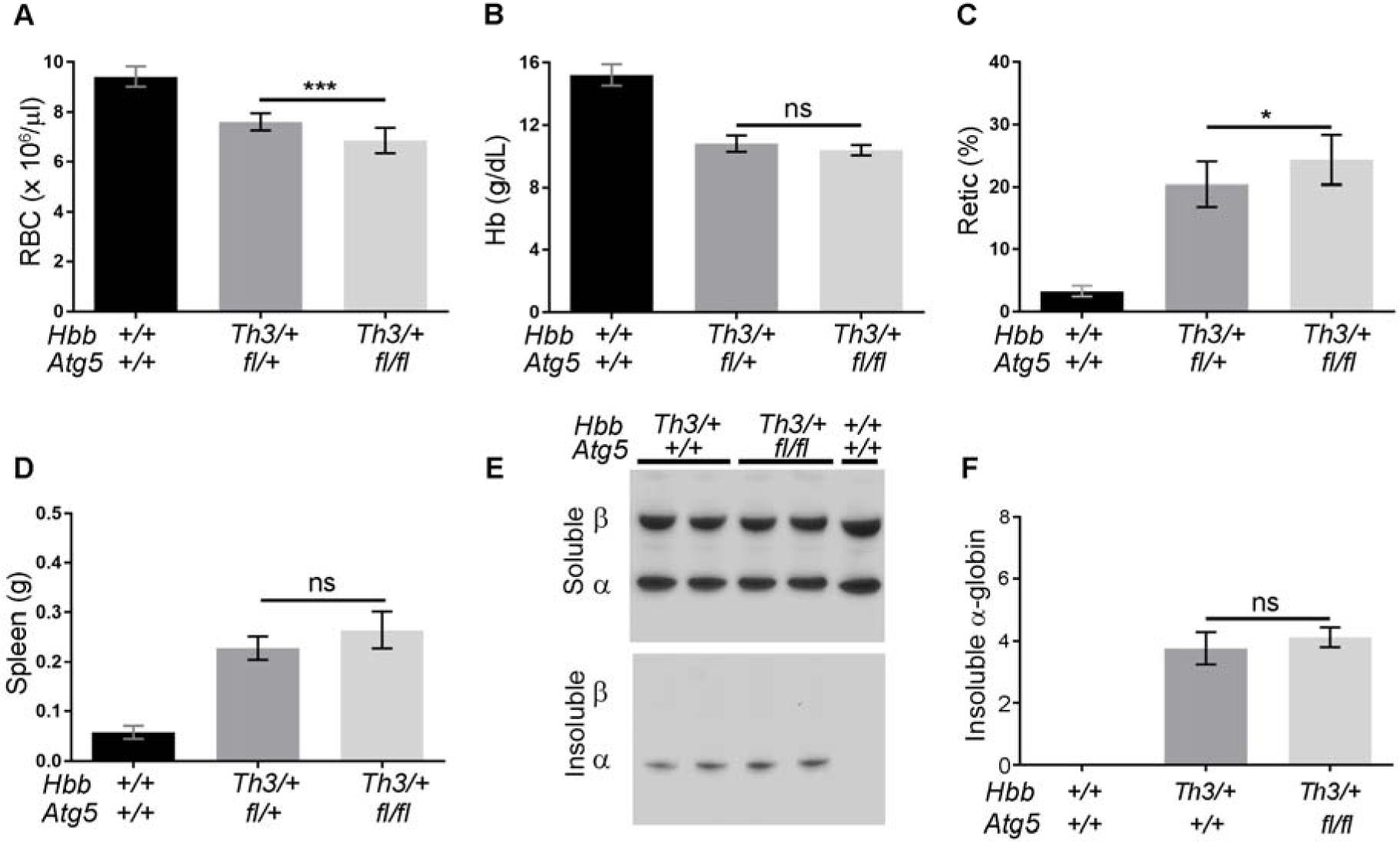
Clearance of free α-globin in β-thalassemia is ATG5-independent. (**A–C**) Erythroid indices (y-axis) of mice with the indicated genotypes (x-axis) at 5 months of age. *Hbb^+/+^Atg5^+/+^*, n = 10; *Hbb*^*Th3*/+^*Atg5*^+/+^, n = 10; *Hbb*^*Th3*/+^*Atg5*^*fl/fl*^, n = 12. (**D**) Spleen weight according to genotype. WT, n = 10; *Hbb*^*Th3*/+^*Atg5*^+/+^, n = 5; *Hbb*^*Th3*/+^*Atg5*^*fl/fl*^, n = 4. (**E**) Soluble and insoluble α-globin in RBCs. Globin chains were resolved by TAU gel electrophoresis, then stained with Coomassie brilliant blue. (**F)** Results of multiple experiments performed as in panel E. The y-axis shows the relative globin staining intensity on TAU gels, measured by automated image analysis (AlphaView SA). WT, *Hbb^+/+^Ulk1^+/+^*, n = 4; *Hbb*^*Th3*/+^*Atg5*^+/+^, n = 4; *Hbb*^*Th3*/+^*Atg5*^*fl/fl*^, n = 4. All graphs show the mean ± SD; \*\*\**P* < 0.001; \**P* < 0.05; ns: not significant.

### Lysosomal clearance of free α-globin in *Hbb*^*Th3*/+^ β-thalassemic mice is ATG5-independent and ULK1-dependent

Previous work by our group(*9*) and others (reviewed in (*11*)) demonstrated that the ubiquitin-proteasome system and autophagy degrade excess α-globin in β-thalassemic erythroid cells. Systemic administration of the proteasome inhibitor bortezomib to *Hbb*^*Th3*/+^ mice induced both autophagy and heat-shock proteins with no net increase in insoluble α-globin(*9*), suggesting that there is cross-compensation by the PQC systems. To investigate how the depletion of ULK1 or ATG5 affects the turnover of insoluble α-globin in β-thalassemia, we pulse-labeled mutant reticulocytes with ^35^S-labeled amino acids, chased for various times with unlabeled amino acids with or without various inhibitors, and quantified the soluble and insoluble nascent radiolabeled α-globin in cell lysates. After 6 h, *Hbb*^Th3/+^ reticulocytes had cleared approximately 60% of the labeled insoluble α-globin through mechanisms that were sensitive to proteasome inhibition with MG132 (MG) or to late stage autophagy inhibition by chloroquine (CQ), or bafilomycin A1 (Baf) (Fig. 4A,B and Fig. S3). In contrast, autophagy-dependent clearance of insoluble α-globin was fully eliminated in *Hbb*^*Th3*/+^*Ulk1*^−/−^ reticulocytes (Fig. 4C,D) but was maintained in *Hbb*^*Th3*/+^*Atg5*^*fl/fl*^ reticulocytes (Fig. 4E,F). Thus, in β-thalassemia, insoluble free α-globin is cleared by autophagy that is ULK1-dependent and ATG5-independent.

**Fig. 4.**
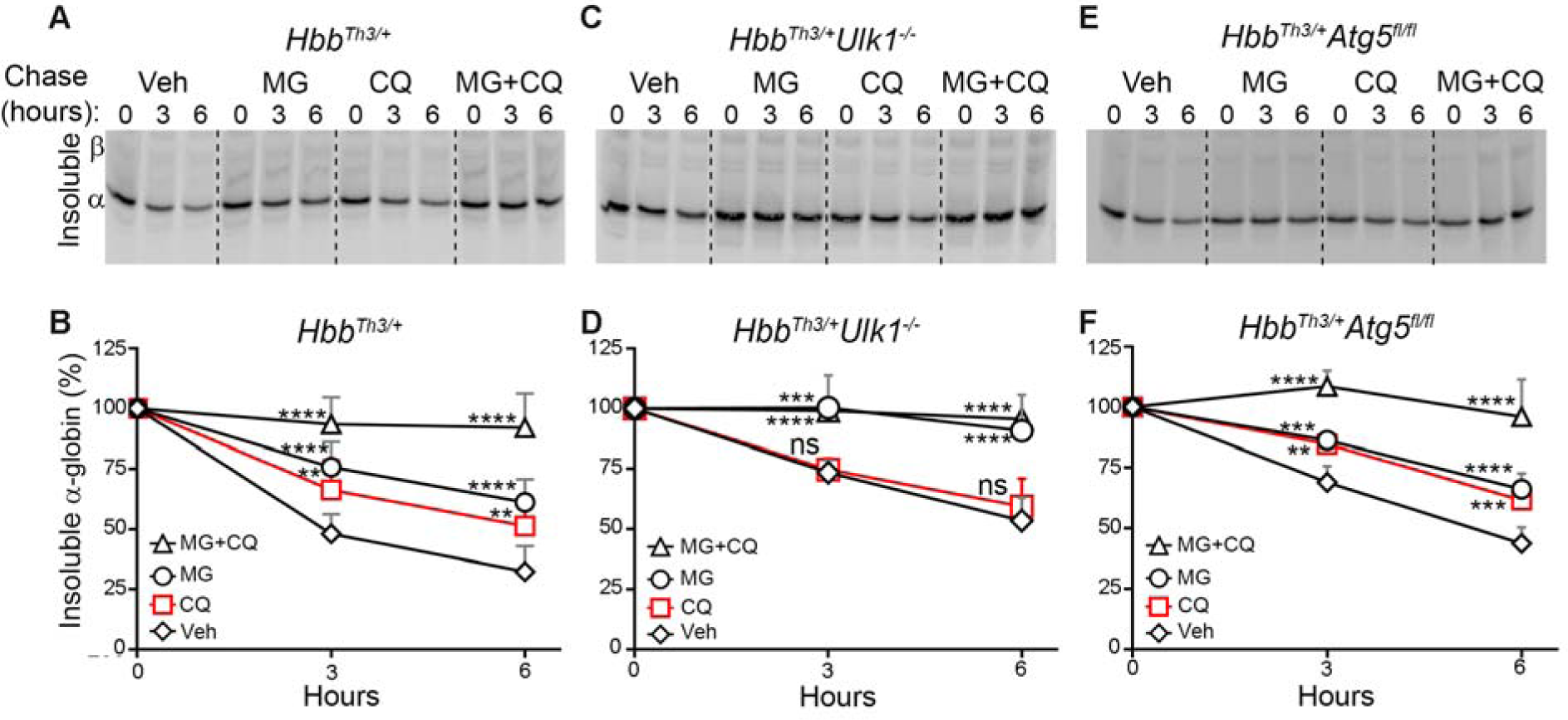
Lysosome-dependent flux of insoluble free α-globin in reticulocytes is Atg-5-independent and Ulk-1-dependent. (**A,C,E**) Reticulocytes in 60μL of whole blood from mice with the indicated genotypes were pulse-labeled with ^35^S-amino acids, then chased for 0, 3, and 6 h with unlabeled amino acids ± proteasome inhibitor (MG132 [MG], 10µM) and/or lysosome inhibitor (chloroquine [CQ], 100µM). Insoluble α-globin in cell lysates was resolved by differential centrifugation and TAU gel electrophoresis (see Fig. 2A) then visualized by autoradiography. (**B,D,F**) Results of multiple experiments performed as in panels a, c, and e. The y-axis shows the relative globin staining intensity on TAU gels as measured by autoradiography and automated image analysis (AlphaView SA). *Hbb*^*Th3*/+^, n = 8; *Hbb*^*Th3*/+^*Ulk1*^−/−^, n = 5; *Hbb*^*Th3*/+^*Atg5*^*fl/fl*^, n = 4. All graphs show the mean ± SD; \*\*\*\**P* < 0.0001; \*\*\**P* < 0.001; \*\**P* < 0.01; ns: not significant versus vehicle for each genotype, respectively, by two-way ANOVA.

### Rapamycin increases free α-globin clearance and reduces pathologies in β-thalassemia via ULK1 activation

mTORC1 inhibits autophagy by phosphorylating ULK1(*42–45*). Therefore, we tested whether mTORC1 inhibition could ameliorate β-thalassemia by stimulating autophagic clearance of free α-globin. Rapamycin (4 mg/kg) was administered intraperitoneally to β-thalassemic and control mice daily for 30 days, followed by necropsy. In vivo drug activity in reticulocytes was confirmed by reduced phosphorylation of the mTORC1 target ribosomal protein S6 (*P* < 0.001) (Fig. S4A,B). β-Thalassemic mice treated with rapamycin exhibited numerous indications of reduced RBC pathology, including a 27% increase in RBC count (*P* < 0.001), a 10% increase in Hb (*P* < 0.05), a 44% decrease in reticulocyte count (*P* < 0.001), and an increase in the half-life of circulating RBCs from 6.9 to 12.2 days (*P* < 0.0001; the normal half-life is 18 days) (Fig. 5A–E and Table S4). Rapamycin-treated *Hbb*^*Th3*/+^ mice also showed signs of diminished ineffective erythropoiesis, including a reduction in spleen weight (*P* < 0.01), as well as reduced levels of bone marrow erythroid hyperplasia and extramedullary erythropoiesis in the spleen and liver (Fig. 5F,G and Table S5). Importantly, rapamycin had no effect on RBC indices or erythropoiesis in WT or *Hbb*^*Th3*/+^*Ulk1*^−/−^ mice.

**Fig. 5.**
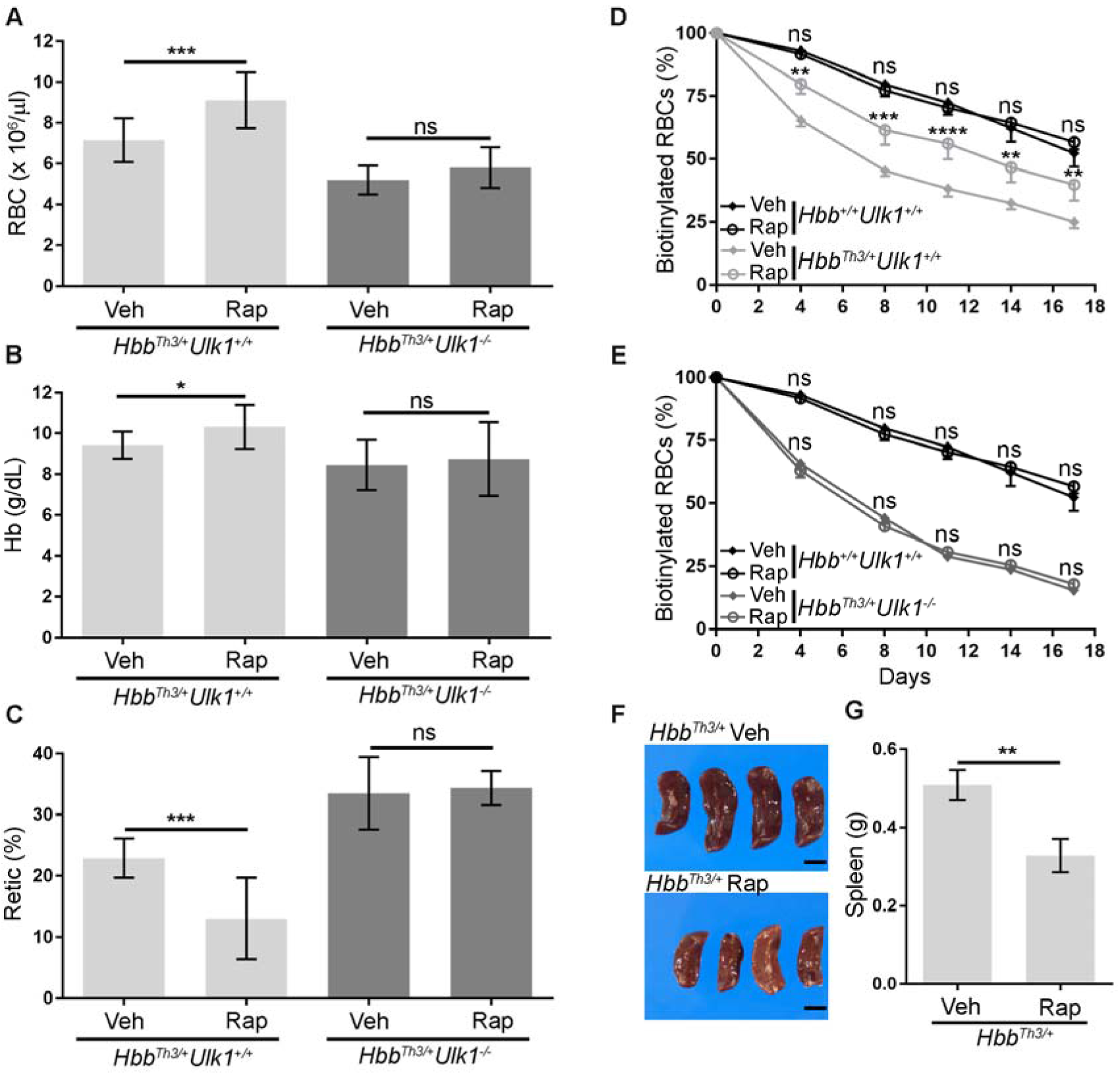
Rapamycin induces ULK1-dependent improved clinical phenotypes in β-thalassemic mice. Wild-type mice were transplanted with fetal liver *Hbb*^*Th3*/+^*Ulk1*^−/−^ HSCs as in Fig. 1. Twelve weeks later, rapamycin (Rap, 4 mg/kg) or vehicle (Veh, 0.5% ethanol, 5% Tween 80, 5% polyethylene glycol 400) were administered intraperitoneally daily for 1 month. (**A–C**) Erythroid indices (y-axis) according to genotype (x-axis) after treatment with rapamycin or vehicle. *Hbb*^*Th3*/+^*Ulk1*^+/+^, n = 12; *Hbb*^*Th3*/+^*Ulk1*^−/−^, n = 9. (**D,E**) RBC survival studies. *Hbb*^*Th3*/+^*Ulk1*^+/+^ mice (n = 7) (**D**) and *Hbb*^*Th3*/+^*Ulk1*^−/−^ mice (n = 4) (**E**) were treated with rapamycin or vehicle for 30 days and injected at day 13 with sulfo-NHS-biotin. The fraction of biotinylated RBCs over time was quantified by streptavidin labeling and flow cytometry. The measured RBC half-lives (in days) were: 18.0 (Veh) and 18.2 (Rap) for *Hbb^+/+^Ulk1^+/+^* (n = 8); 6.9 (Veh) and 12.2 (Rap) for *Hbb*^*Th3*/+^*Ulk1*^+/+^ (n = 7); and 6.1 (Veh) and 6.2 (Rap) days for *Hbb*^*Th3*/+^*Ulk1*^−/−^ (n = 4). (**F**) Spleens of β-thalassemic mice (*Hbb*^*Th3*/+^) after rapamycin (Rap) or vehicle treatment for 30 days. (**G**) Summary of spleen weights in multiple *Hbb*^*Th3*/+^ mice after treatment with vehicle (n = 9) or rapamycin (n = 8). All graph shows the mean ± SD; \*\*\*\**P* < 0.0001; \*\*\**P* < 0.001; \*\**P* < 0.01; \**P* < 0.05; ns: not significant.

We studied RBCs and their precursors to investigate the mechanisms by which rapamycin ameliorates the effects of β-thalassemia. Insoluble α-globin in reticulocytes from rapamycin-treated *Hbb*^*Th3*/+^ mice was decreased by approximately 50% (*P* < 0.01) (Fig. 6A–D). Consistent with these findings, TEM showed an approximately 60% reduction in electron-dense aggregates in reticulocytes from β-thalassemic mice treated with rapamycin, as compared to those from vehicle-treated controls (*P* < 0.0001) (Fig. 6E,F). The levels of autophagosomal membrane-bound LC3-II decreased in reticulocytes of *Hbb*^*Th3*/+^ mice treated with rapamycin (Fig. 6G), suggesting enhanced flux through the autophagy pathway in these animals. In contrast, rapamycin had no effect on the accumulation of insoluble α-globin in *Hbb*^*Th3*/+^*Ulk1*^−/−^ mice (Fig. 6C–G). Together, these studies demonstrate that rapamycin treatment of β-thalassemic mice stimulates ULK1-dependent autophagy to reduce free α-globin precipitates, alleviate ineffective erythropoiesis, and extend RBC lifespan.

**Fig. 6.**
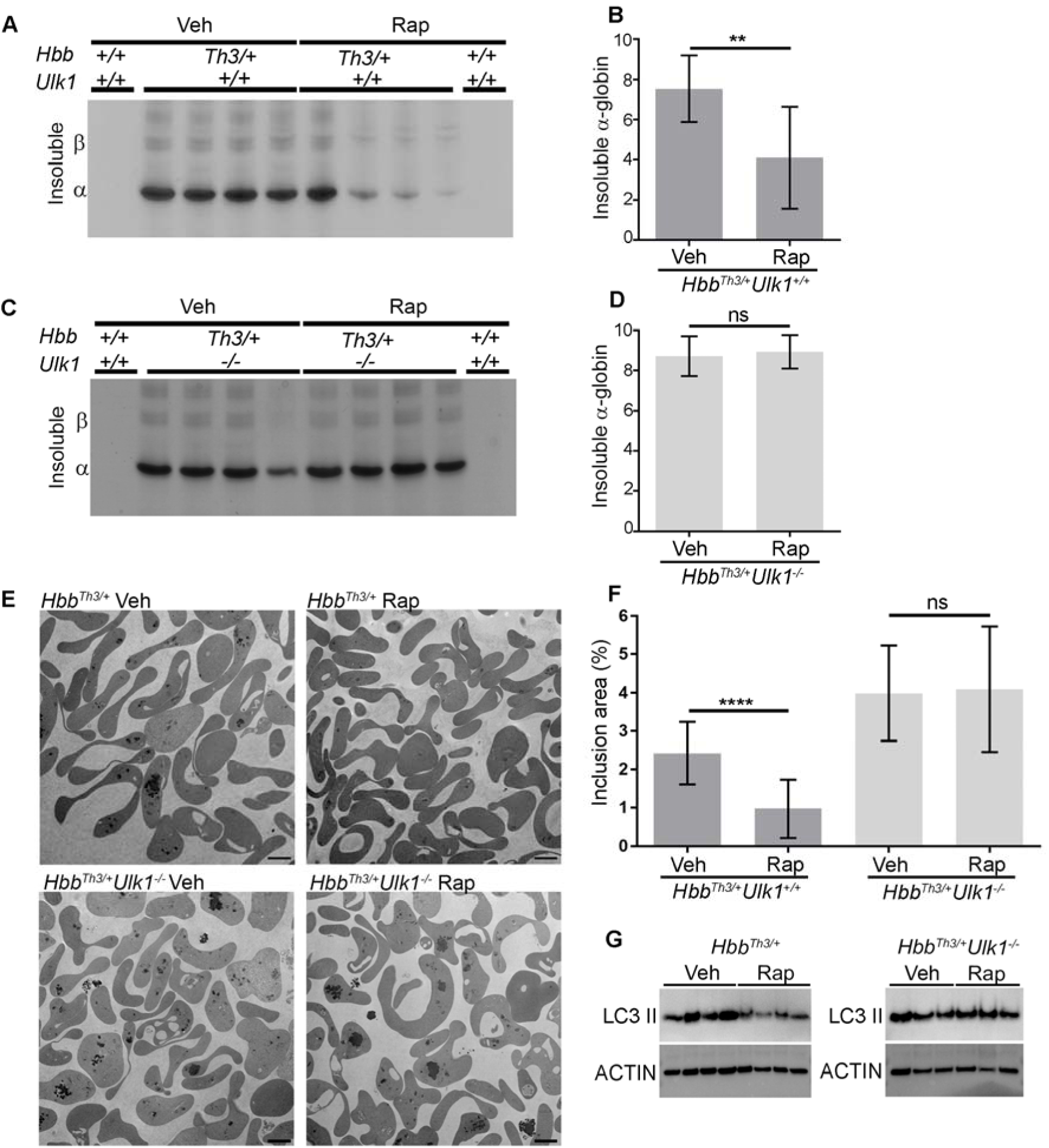
Rapamycin induces ULK1-dependent elimination of free α-globin. (**A,C**) Insoluble α-globin levels after 30-day treatment with rapamycin (Rap) or vehicle (Veh) of mice transplanted with *Hbb*^*Th3*/+^*Ulk1*^+/+^ or *Hbb*^*Th3*/+^*Ulk1*^−/−^ HSCs. RBCs were lysed and analyzed as described for Fig. 2C. (**B,D**) Results of multiple experiments performed as in panels a and c. The y-axis shows the relative α-globin staining intensity on TAU gels as measured by automated image analysis. *Hbb*^*Th3*/+^ + Veh, n = 7; *Hbb*^*Th3*/+^ + Rap n = 7; *Hbb*^*Th3*/+^*Ulk1*^−/−^ + Veh, n = 7; *Hbb*^*Th3*/+^*Ulk1*^−/−^ + Rap, n = 7. (**E**) Electron micrographs of purified reticulocytes in mice of the indicated genotypes treated with Rap or Veh. Scale bars represent 2 µm. (**F**) Areas of electron-dense α-globin inclusions in reticulocytes (y-axis) determined by automated image analysis of electron micrographs *Hbb*^*Th3*/+^*Ulk1*^+/+^ + Veh, n = 8; *Hbb*^*Th3*/+^*Ulk1*^+/+^ + Rap, n = 7; *Hbb*^*Th3*/+^*Ulk1*^−/−^ + Veh, n = 7; *Hbb*^*Th3*/+^*Ulk1*^−/−^ + Rap, n = 7. (**G**) Western blot analysis for autophagic markers of reticulocytes from mice of the indicated genotypes treated with Rap or Veh. All graphs show the mean ± SD; \*\*\*\**P* < 0.0001; \*\**P* < 0.01; ns: not significant.

### Rapamycin reduces free α-globin in human β-thalassemic erythroblasts

To investigate the therapeutic potential of rapamycin in human β-thalassemia, we treated erythroid precursors generated by *in vitro* differentiation of CD34^+^ hematopoietic stem and progenitor cells (HSPCs) from patients with β-thalassemia who require regular RBC transfusions (transfusion-dependent, TD) (n = 5) and from those with variable degrees of anemia who require transfusions intermittently or not at all (non–transfusion-dependent, NTD) (n = 5) (Table S6 and Table S7). Patient or control CD34^+^ cells were purified from either peripheral blood (for control and NTD β-thalassemia cells) or bone marrow (for TD β-thalassemia cells) and grown in culture under conditions that support erythroid differentiation(*46*). β-Thalassemia erythroblasts exhibited accelerated maturation, as indicated by the earlier appearance of the late-stage erythroid marker Band3 and the loss of the integrin subunit CD49d (Fig. 7A,B and Fig. S5A,B). Reverse-phase high-performance liquid chromatography (RP-HPLC) analysis of hemoglobinized erythroblasts generated from patients with TD or NTD β-thalassemia revealed α-chain excesses (α-chain/β-like chain [β+γ+δ]) of approximately 40% and 15%, respectively (Fig. S5C,D). Non-denaturing ion-exchange (IE) HPLC identified a free α-globin peak in TD and NTD β-thalassemic erythroblasts, but not in control cells (Fig. 7C–F and Fig. S5E,F). Rapamycin (10µM or 20µM) or the proteasome inhibitor MG132 was added to day 13 cultures, which contained mid- to late-stage erythroblasts, and α-globin accumulation was determined by HPLC 2 days later. As expected, proteasome inhibition by MG132 raised free α-globin levels in β-thalassemic erythroblasts and also induced cell death (Fig. 7C–F, and Fig. S6A–D). In contrast, rapamycin treatment reduced free α-globin by 40% and 85% in TD β-thalassemia (*P* < 0.0001) and NTD β-thalassemic erythroblasts (*P* < 0.001), respectively, with no deleterious effect on cell survival (Fig. 7C–F and Fig. S6A–D). Previous studies indicated that rapamycin could elevate fetal hemoglobin level in cultures of erythroblasts derived from human CD34^+^ cells(*47, 48*), even if we did not observe this effect in our studies (Fig. S6E). Thus, rapamycin reduces the accumulation of free α-globin in primary erythroid cells derived from patients with β-thalassemia, most likely by inhibiting mTORC1, with consequent derepression of ULK1-mediated autophagy.

**Fig. 7.**
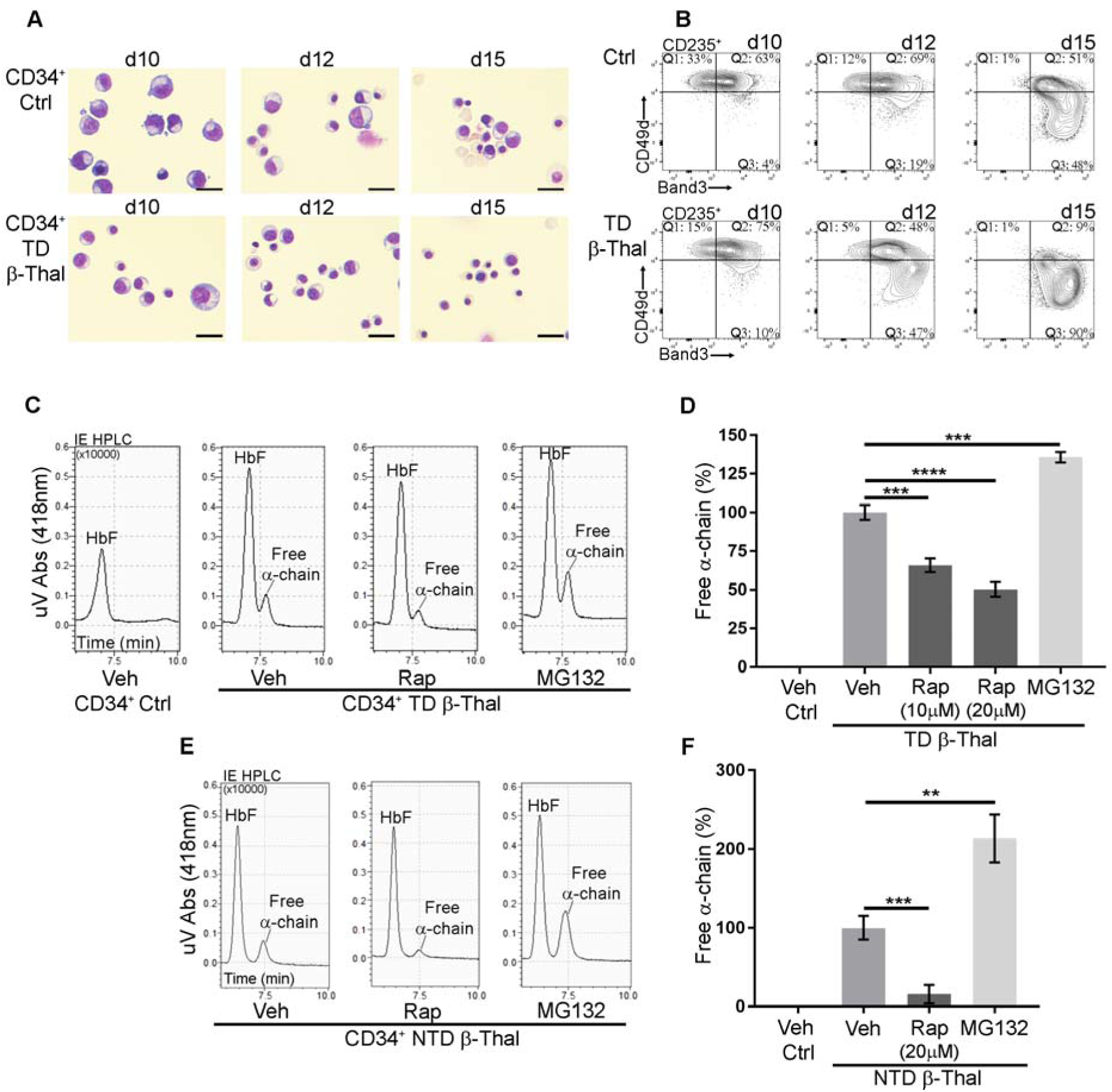
Rapamycin reduces insoluble α-globin in erythroblasts derived from CD34^+^ cells of human patients with β-thalassemia. Peripheral blood CD34^+^ cells from normal donors or from patients with β-thalassemia were grown in culture to promote erythroid differentiation. (**A**) May-Grunwald–stained cytocentrifugation preparations on the indicated days of erythroid culture. Scale bars represent 10µm. (**B**) Representative flow cytometry plots showing the expression of erythroid maturation markers (CD235 and Band3) and the immature cell marker CD49d at the indicated days of culture. (**C,E**) CD34^+^ cell-derived erythroblasts from patients with transfusion-dependent (TD) (**C**) or non–transfusion-dependent (NTD) (**E**) β-thalassemia were treated with rapamycin (10µM or 20µM) or MG132 (2.5µM) on culture days 13–15 then analyzed for free α-globin by isoelectric focusing (IE)-HPLC. Representative chromatographs are shown, with free α-globin indicated. (**D,F**) Summary of multiple experiments performed as in panels c and e. Free α-chain was quantified by image analysis of HPLC data (from seven healthy individuals, five individuals with TD β-thalassemia, and five individuals with NTD β-thalassemia). All graphs show the mean ± SD; \*\*\*\**P* < 0.0001; \*\*\**P* < 0.001; \*\**P* < 0.01

### Discussion

We show that ULK1-dependent autophagy of free α-globin reduces ineffective erythropoiesis and improves RBC viability in β-thalassemia and that this process can be stimulated by the mTORC1 inhibitor rapamycin in mice and in human erythroid cells. Our findings are consistent with the longstanding concepts that globin protein imbalance with excessive build-up of toxic free α-chain is a major determinant of β-thalassemia severity(*49*) and that β-thalassemia can be classified as a “protein aggregation disorder” in which disease severity is modulated by PQC activities including the ubiquitin-proteasome system and autophagy^(*11*)^. The identification of ULK1 as a central regulator of free α-globin autophagy elucidates further our understanding of β-thalassemia and highlights a “druggable” pathway that can be leveraged therapeutically.

Free α-globin clearance in erythroid cells is summarized in our current model. In β-thalassemia, unpaired α-chains are polyubiquitinated and subsequently degraded by the proteasome. The erythroid-specific ubiquitin ligase UBE20 ubiquitinates free α-globin (*50, 51*), although additional ubiquitin ligases likely exist. Alternatively, free α-globin (with or without a ubiquitin modification) may be eliminated by autophagy. In β-thalassemic mice, autophagy-mediated degradation of free α-globin is stimulated by proteasome inhibition^9^, which is consistent with cross-regulation between the UPS and autophagy pathways(*9, 52*). In contrast, our current findings show that the UPS does not compensate for the loss of ULK1-mediated α-globin autophagy, which is consistent with studies demonstrating that autophagy inhibition impairs the degradation of proteasome substrates(*53*).

Autophagy can occur via multiple mechanisms depending on the physiologic context, cell type, and cargo, with substantial variation in the requirements for specific autophagy proteins (reviewed in reference (*24*)). Autophagy of free α-globin requires ULK1, but not the canonical autophagy protein ATG5, which facilitates the conjugation of LC3 and related proteins to the autophagosome outer membrane(*54*). ULK1-dependent, ATG5-independent autophagy also occurs after DNA damage^58^, and is required for the efficient elimination of mitochondria during reprogramming of fibroblasts to induced pluripotent stem cells(*55*) and maturation of fetal reticulocytes(*37, 56*),^59^. In the previous studies, ATG5-independent autophagy required the GTPase RAB9, probably for the trafficking of golgi vesicles to elongating autophagosomes. Further studies will be important to better understand the mechanisms of free α-globin autophagy. Of particular interest will be to define the requirements for various autophagy-related proteins including RAB9, and to elucidate the mechanisms that direct free α-globin to autophagosomes.

ULK1 is positively regulated by phosphorylation from AMPK and is inhibited by mTORC1-mediated phosphorylation^(*45, 57, 58*)^. We tested rapamycin as an ULK1 activator in β-thalassemia because it is an approved drug with a well-defined, manageable toxicity profile(*47*). Rapamycin or related drugs stimulate neuronal autophagy of pathological proteins in several experimental models for neurodegenerative disease(*59*) and improve RBC number in β-thalassemic mice(*60*). Our findings extend the latter result by elucidating the definitive mechanism (ULK1-dependent autophagy of free α-globin) and showing that rapamycin can reduce free α-globin accumulation in human erythroblasts. Specially, rapamycin treatment significantly reduced erythroblast accumulation of free α-globin in a dose-dependent manner in TD β-thalassemic cells and almost fully eliminated free α-globin in NTD β-thalassemic cells treated with the highest dose of rapamycin tested. In addition to stimulating autophagy, rapamycin treatment of cultured human erythroblasts induces the synthesis of γ-globin, a fetal-expressed β-like globin that can bind and detoxify free α-globin(*47*). While this effect did not occur in our studies, probably due to differences in experimental design, rapamycin-induced γ-globin synthesis could synergize with enhanced α-globin autophagy to further alleviate the pathophysiology of β-thalassemia. Of note, this hypothesis cannot be tested easily in mouse models for β-thalassemia because fetal-expressed γ-globin genes exist only in humans and some nonhuman primates. Overall, we believe that the composite data now provide sufficient rationale for a clinical trial to test the effect of treatment with rapamycin or a newer analog on ineffective erythropoiesis and RBC longevity in β-thalassemia. Moreover, the identification of ULK1 as a major effector of clearance of free α-globin identifies new therapeutic targets, namely AMPK and ULK1 itself. The development of drugs that activate these kinases is underway(*61, 62*), and their utility as potential therapies for β-thalassemia should be assessed. More generally, our findings should arouse new interest in the concept of therapeutic ULK activation to stimulate autophagy in other protein aggregation disorders.

## Acknowledgments

We are grateful to Sharon Frase, Linda Horner, and Richard Gursky (St. Jude Cellular Imaging Shared Research Core); Sean Savage, Michael Anderson, Patty Varner, Brenda Gallini, and Michael Straign (St. Jude Veterinary and Pathology Core); Dick Ashmun and all the staff (Flow Cytometry and Cell Sorting Shared Resource); Shandra Savage, Madoka Inoue, and all the staff (St. Jude Animal Resource Center); Valentina Brancaleoni (University of Milan, Genotyping Resource Center). We thank Doug Green for helpful discussions. We thank Anita Impagliazzo for her graphic design of our model. We thank Keith A. Laycock (St. Jude Department of Scientific Editing) for editing the manuscript.

## Author contributions

C.L. designed and performed experiments, analyzed and interpreted data, and wrote the paper and edited the manuscript. J.K. performed experiments, assisted in data analysis and edited the manuscript. E.K. designed and performed experiments, and analyzed data on *Hbb*^*Th3*/+^*Atg5*^*fl/fl*^ mice. S.F., K.M., A.F., C.S.T, P.D, E.B.R., and J.Z. performed experiments and/or assisted in data analysis. I.M and M.D.C. provided human CD34^+^ cells and clinical data, and edited the manuscript. H.T. performed experiments, analyzed physiopathology data, and assisted in automated image analysis of electron micrographs. M.K. conceived the project, provided *Ulk1^−/−^* mice and edited the manuscript. M.J.W. conceived the project, interpreted data and wrote and edited the manuscript.

## Data and materials availability

All data are available in the main text or the supplementary materials.

## Supplementary Materials

Materials and Methods

Fig. S1 to S6

Tables S1 to S7

References

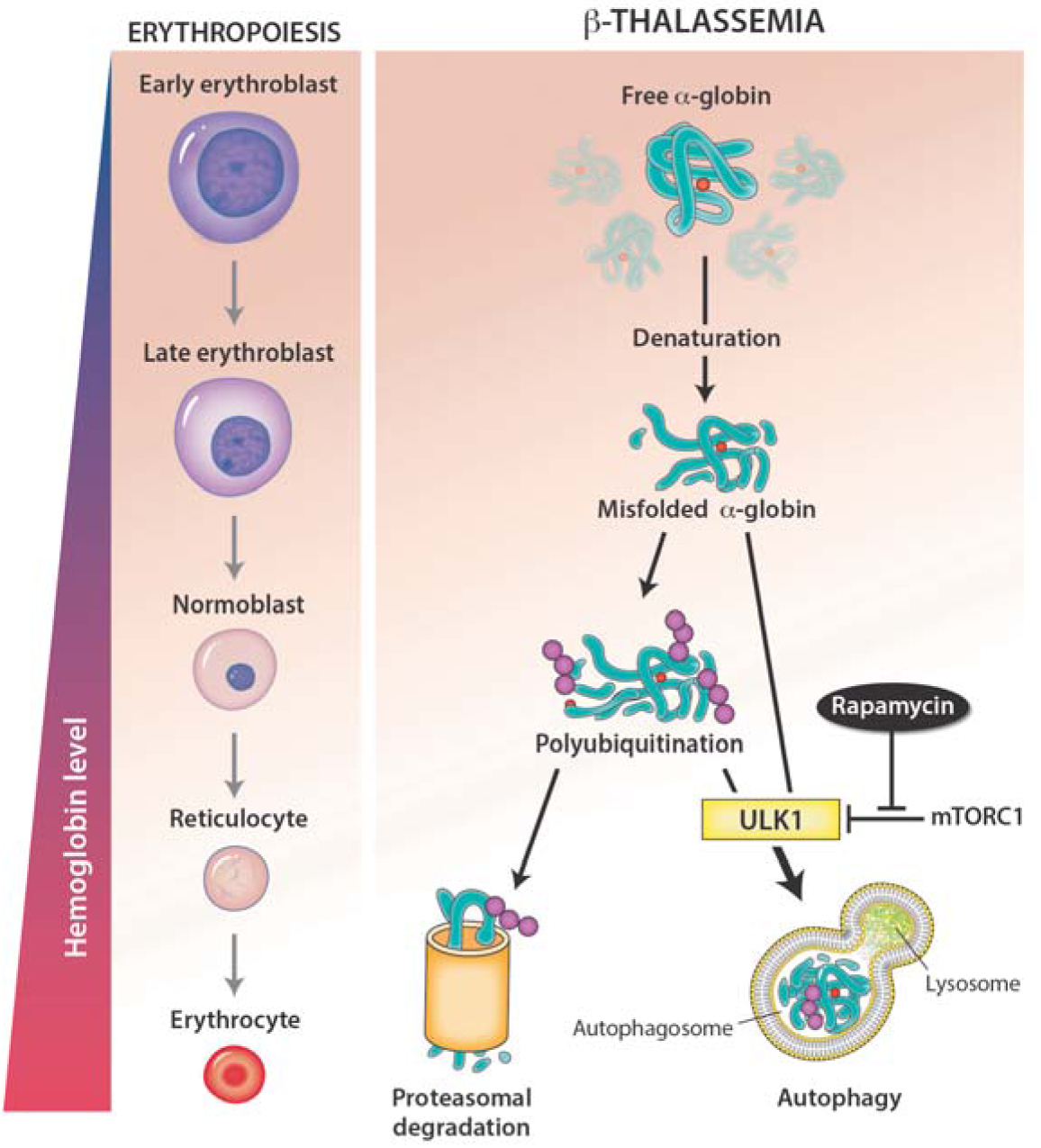
Model for ULK1 kinase-dependent clearance of free α-globin in β-thalassemia. During mammalian erythroblast differentiation, the hemoglobin concentration gradually increases. In β-thalassemia, β-globin gene mutations cause a build-up of free α-globin, which forms intracellular precipitates that impair erythroid cell maturation and viability. Free α-globin is eliminated by protein quality-control pathways, including the ubiquitinproteasome system and autophagy. Rapamycin treatment significantly reduced the accumulation of free α-globin in an *Ulk1*-dependent fashion.

## References

1. D. R. Higgs, J. D. Engel, G. Stamatoyannopoulos, Thalassaemia. Lancet 379, 373–383 (2012).

2. M. H. Steinberg, Disorders of hemoglobin : genetics, pathophysiology, and clinical management. (Cambridge University Press, Cambridge, ed. 2nd ed., 2009), pp. xx, 826 p., [836] p. of plates.

3. V. Vigi et al., The correlation between red-cell survival and excess of alpha-globin synthesis in beta-thalassemia. Br J Haematol 16, 25–30 (1969).

4. E. Shinar, O. Shalev, E. A. Rachmilewitz, S. L. Schrier, Erythrocyte membrane skeleton abnormalities in severe beta-thalassemia. Blood 70, 158–164 (1987).

5. R. Advani, E. Rubin, N. Mohandas, S. L. Schrier, Oxidative red blood cell membrane injury in the pathophysiology of severe mouse beta-thalassemia. Blood 79, 1064–1067 (1992).

6. E. Rachmilewitz, S. Schrier, Pathophysiology of Beta Thalassemia. Disorders of Hemoglobin: Genetics, (2001).

7. S. H. Orkin et al., Nathan and Oski’s hematology of infancy and childhood. (Saunders, Philadelphia, Pa., ed. 7th, 2008).

8. M. C. Sollaino et al., Association of alpha globin gene quadruplication and heterozygous beta thalassemia in patients with thalassemia intermedia. Haematologica 94, 1445–1448 (2009).

9. E. Khandros, C. S. Thom, J. D’Souza, M. J. Weiss, Integrated protein quality-control pathways regulate free alpha-globin in murine beta-thalassemia. Blood 119, 5265–5275 (2012).

10. A. Premawardhena et al., A novel molecular basis for beta thalassemia intermedia poses new questions about its pathophysiology. Blood 106, 3251–3255 (2005).

11. E. Khandros, M. J. Weiss, Protein quality control during erythropoiesis and hemoglobin synthesis. Hematol Oncol Clin North Am 24, 1071–1088 (2010).

12. D. R. Higgs, The thalassaemia syndromes. Q J Med 86, 559–564 (1993).

13. H. Birgens, R. Ljung, The thalassaemia syndromes. Scand J Clin Lab Invest 67, 11–25 (2007).

14. Y. Kong et al., Loss of alpha-hemoglobin-stabilizing protein impairs erythropoiesis and exacerbates beta-thalassemia. The Journal of clinical investigation 114, 1457–1466 (2004).

15. A. J. Kihm et al., An abundant erythroid protein that stabilizes free alpha-haemoglobin. Nature 417, 758–763 (2002).

16. A. Aigelsreiter et al., How a cell deals with abnormal proteins. Pathogenetic mechanisms in protein aggregation diseases. Pathobiology 74, 145–158 (2007).

17. A. Ciechanover, P. Brundin, The ubiquitin proteasome system in neurodegenerative diseases: sometimes the chicken, sometimes the egg. Neuron 40, 427–446 (2003).

18. S. C. Meredith, Protein denaturation and aggregation: Cellular responses to denatured and aggregated proteins. Ann N Y Acad Sci 1066, 181–221 (2005).

19. D. Cox, C. Raeburn, X. Sui, D. M. Hatters, Protein aggregation in cell biology: An aggregomics perspective of health and disease. Semin Cell Dev Biol, (2018).

20. P. Lithanatudom et al., Enhanced activation of autophagy in beta-thalassemia/Hb E erythroblasts during erythropoiesis. Ann Hematol 90, 747–758 (2011).

21. N. Mizushima, T. Yoshimori, Y. Ohsumi, The role of Atg proteins in autophagosome formation. Annu Rev Cell Dev Biol 27, 107–132 (2011).

22. M. S. Uddin et al., Autophagy and Alzheimer’s Disease: From Molecular Mechanisms to Therapeutic Implications. Front Aging Neurosci 10, 04 (2018).

23. A. Ciechanover, Y. T. Kwon, Degradation of misfolded proteins in neurodegenerative diseases: therapeutic targets and strategies. Exp Mol Med 47, e147 (2015).

24. L. Galluzzi et al., Molecular definitions of autophagy and related processes. EMBO J 36, 1811–1836 (2017).

25. E. Masiero et al., Autophagy is required to maintain muscle mass. Cell metabolism 10, 507–515 (2009).

26. T. Hara et al., Suppression of basal autophagy in neural cells causes neurodegenerative disease in mice. Nature 441, 885–889 (2006).

27. A. Nakai et al., The role of autophagy in cardiomyocytes in the basal state and in response to hemodynamic stress. Nature medicine 13, 619–624 (2007).

28. M. Komatsu, Y. Ichimura, Selective autophagy regulates various cellular functions. Genes Cells 15, 923–933 (2010).

29. N. Mizushima, A brief history of autophagy from cell biology to physiology and disease. Nat Cell Biol 20, 521–527 (2018).

30. P. Codogno, M. Mehrpour, T. Proikas-Cezanne, Canonical and non-canonical autophagy: variations on a common theme of self-eating? Nature reviews. Molecular cell biology 13, 7–12 (2011).

31. N. T. Ktistakis, S. A. Tooze, Digesting the Expanding Mechanisms of Autophagy. Trends Cell Biol 26, 624–635 (2016).

32. L. H. Travassos, L. R. Vasconcellos, M. T. Bozza, L. A. Carneiro, Heme and iron induce protein aggregation. Autophagy 13, 625–626 (2017).

33. B. Wang, M. Kundu, Canonical and noncanonical functions of ULK/Atg1. Curr Opin Cell Biol 45, 47–54 (2017).

34. N. N. Noda, Y. Fujioka, Atg1 family kinases in autophagy initiation. Cell Mol Life Sci 72, 3083–3096 (2015).

35. M. Kundu et al., Ulk1 plays a critical role in the autophagic clearance of mitochondria and ribosomes during reticulocyte maturation. Blood 112, 1493–1502 (2008).

36. L. S. Tain et al., Rapamycin activation of 4E-BP prevents parkinsonian dopaminergic neuron loss. Nat Neurosci 12, 1129–1135 (2009).

37. S. Honda et al., Ulk1-mediated Atg5-independent macroautophagy mediates elimination of mitochondria from embryonic reticulocytes. Nat Commun 5, 4004 (2014).

38. M. Kundu, ULK1, mammalian target of rapamycin, and mitochondria: linking nutrient availability and autophagy. Antioxid Redox Signal 14, 1953–1958 (2011).

39. B. Yang et al., A mouse model for beta 0-thalassemia. Proceedings of the National Academy of Sciences of the United States of America 92, 11608–11612 (1995).

40. A. C. Heinrich, R. Pelanda, U. Klingmuller, A mouse model for visualization and conditional mutations in the erythroid lineage. Blood 104, 659–666 (2004).

41. W. J. Liu et al., p62 links the autophagy pathway and the ubiqutin-proteasome system upon ubiquitinated protein degradation. Cell Mol Biol Lett 21, 29 (2016).

42. M. Laplante, D. M. Sabatini, mTOR signaling in growth control and disease. Cell 149, 274–293 (2012).

43. I. G. Ganley et al., ULK1.ATG13.FIP200 complex mediates mTOR signaling and is essential for autophagy. The Journal of biological chemistry 284, 12297–12305 (2009).

44. N. Hosokawa et al., Nutrient-dependent mTORC1 association with the ULK1-Atg13-FIP200 complex required for autophagy. Mol Biol Cell 20, 1981–1991 (2009).

45. J. Kim, M. Kundu, B. Viollet, K. L. Guan, AMPK and mTOR regulate autophagy through direct phosphorylation of Ulk1. Nat Cell Biol 13, 132–141 (2011).

46. E. A. Traxler et al., A genome-editing strategy to treat beta-hemoglobinopathies that recapitulates a mutation associated with a benign genetic condition. Nature medicine 22, 987–990 (2016).

47. E. Fibach et al., Effects of rapamycin on accumulation of alpha-, beta- and gamma-globin mRNAs in erythroid precursor cells from beta-thalassaemia patients. Eur J Haematol 77, 437–441 (2006).

48. A. Pecoraro et al., Efficacy of Rapamycin as Inducer of Hb F in Primary Erythroid Cultures from Sickle Cell Disease and beta-Thalassemia Patients. Hemoglobin 39, 225–229 (2015).

49. S. Mettananda, R. J. Gibbons, D. R. Higgs, alpha-Globin as a molecular target in the treatment of beta-thalassemia. Blood 125, 3694–3701 (2015).

50. K. Yanagitani, S. Juszkiewicz, R. S. Hegde, UBE2O is a quality control factor for orphans of multiprotein complexes. Science 357, 472–475 (2017).

51. A. T. Nguyen et al., UBE2O remodels the proteome during terminal erythroid differentiation. Science 357, (2017).

52. A. Lilienbaum, Relationship between the proteasomal system and autophagy. Int J Biochem Mol Biol 4, 1–26 (2013).

53. V. I. Korolchuk, A. Mansilla, F. M. Menzies, D. C. Rubinsztein, Autophagy inhibition compromises degradation of ubiquitin-proteasome pathway substrates. Mol Cell 33, 517–527 (2009).

54. J. Geng, D. J. Klionsky, The Atg8 and Atg12 ubiquitin-like conjugation systems in macroautophagy. ‘Protein modifications: beyond the usual suspects’ review series. EMBO Rep 9, 859–864 (2008).

55. T. Ma et al., Atg5-independent autophagy regulates mitochondrial clearance and is essential for iPSC reprogramming. Nat Cell Biol 17, 1379–1387 (2015).

56. Y. Nishida et al., Discovery of Atg5/Atg7-independent alternative macroautophagy. Nature 461, 654–658 (2009).

57. J. W. Lee, S. Park, Y. Takahashi, H. G. Wang, The association of AMPK with ULK1 regulates autophagy. PloS one 5, e15394 (2010).

58. L. Shang et al., Nutrient starvation elicits an acute autophagic response mediated by Ulk1 dephosphorylation and its subsequent dissociation from AMPK. Proceedings of the National Academy of Sciences of the United States of America 108, 4788–4793 (2011).

59. J. Bove, M. Martinez-Vicente, M. Vila, Fighting neurodegeneration with rapamycin: mechanistic insights. Nat Rev Neurosci 12, 437–452 (2011).

60. M. Bayeva et al., mTOR regulates cellular iron homeostasis through tristetraprolin. Cell Metab 16, 645–657 (2012).

61. L. Zhang et al., Discovery of a small molecule targeting ULK1-modulated cell death of triple negative breast cancer in vitro and in vivo. Chem Sci 8, 2687–2701 (2017).

62. J. Li, L. Zhong, F. Wang, H. Zhu, Dissecting the role of AMP-activated protein kinase in human diseases. Acta Pharm Sin B 7, 249–259 (2017).

63. D. J. Ciavatta, T. M. Ryan, S. C. Farmer, T. M. Townes, Mouse model of human beta zero thalassemia: targeted deletion of the mouse beta maj- and beta min-globin genes in embryonic stem cells. Proceedings of the National Academy of Sciences of the United States of America 92, 9259–9263 (1995).

64. K. O. Gudmundsson, S. W. Stull, J. R. Keller, Transplantation of mouse fetal liver cells for analyzing the function of hematopoietic stem and progenitor cells. Methods Mol Biol 879, 123–133 (2012).

65. B. P. Alter, Gel electrophoretic separation of globin chains. Prog Clin Biol Res 60, 157–175 (1981).

66. X. Yu et al., An erythroid chaperone that facilitates folding of alpha-globin subunits for hemoglobin synthesis. The Journal of clinical investigation 117, 1856–1865 (2007).

67. S. Sorensen, E. Rubin, H. Polster, N. Mohandas, S. Schrier, The role of membrane skeletal-associated alpha-globin in the pathophysiology of beta-thalassemia. Blood 75, 1333–1336 (1990)

68. H. Beauchemin, M. J. Blouin, M. Trudel, Differential regulatory and compensatory responses in hematopoiesis/erythropoiesis in alpha- and beta-globin hemizygous mice. The Journal of biological chemistry 279, 19471–19480 (2004).

69. J. Schindelin et al., Fiji: an open-source platform for biological-image analysis. Nat Methods 9, 676–682 (2012).

70. I. Arganda-Carreras et al., Trainable Weka Segmentation: a machine learning tool for microscopy pixel classification. Bioinformatics 33, 2424–2426 (2017).

